# High Throughput Characterization of V(D)J Recombination Signal Sequences Redefines the Consensus Sequence

**DOI:** 10.1101/2021.12.03.471157

**Authors:** Walker Hoolehan, Justin C. Harris, Jennifer N. Byrum, Karla K. Rodgers

**Affiliations:** Department of Biochemistry & Molecular Biology, University of Oklahoma Health Sciences Center, Oklahoma City, Oklahoma

## Abstract

In the adaptive immune system, V(D)J recombination initiates the production of a diverse antigen receptor repertoire in developing B and T cells. Recombination activating proteins, RAG1 and RAG2 (RAG1/2), catalyze V(D)J recombination by cleaving adjacent to recombination signal sequences (RSSs) that flank antigen receptor gene segments. Previous studies defined the consensus RSS as containing conserved heptamer and nonamer sequences separated by a less conserved 12 or 23 base-pair spacer sequence. However, many RSSs deviate from the consensus sequence. Here, we developed a cell-based, massively parallel V(D)J recombination assay to evaluate RAG1/2 activity on thousands of RSSs. We focused our study on the RSS heptamer and adjoining spacer region, as this region undergoes extensive conformational changes during RAG-mediated DNA cleavage. While the consensus heptamer sequence (CACAGTG) was marginally preferred, RAG1/2 was highly active on a wide range of non-consensus sequences. RAG1/2 generally preferred select purine/pyrimidine motifs that may accommodate heptamer unwinding in the RAG1/2 active site. Our results suggest RAG1/2 specificity for RSS heptamers is primarily dictated by DNA structural features dependent on purine/pyrimidine pattern, and to a lesser extent, RAG:RSS base-specific interactions. Further investigation of RAG1/2 specificity using this new approach will help elucidate the genetic instructions guiding V(D)J recombination.

**Summary Statement:** Partially conserved recombination signal sequences (RSSs) govern antigen receptor gene assembly during V(D)J recombination. Here, a massively parallel analysis of randomized RSSs reveals key attributes that allow DNA sequence diversity in the RAG1/2 active site and that contribute to the differential utilization of RSSs in endogenous V(D)J recombination. Overall, these results will assist identification of RAG1/2 off-target sites, which can drive leukemia cell transformation, as well as characterization of *bona fide* RSSs used to generate antigen receptor diversity.

## INTRODUCTION

Functional immunoglobulin and T cell Receptor (TCR) genes are generated in developing B and T cells, respectively, through multiple V(D)J recombination reactions (1,2). This process leads to a diverse repertoire of antigen receptors (AgR) in the adaptive immune system, which are capable of recognizing a vast array of foreign antigens. V(D)J recombination joins V, D, and J gene (coding) segments to construct AgR genes through a cut-and-paste mechanism. V(D)J recombinase specificity guides AgR gene assembly by dictating gene segment utilization in developing B- and T-cells. Gene segments within the same class, such as V_H_ gene segments, are not equally utilized during V(D)J recombination, resulting in biased AgR expression (3).

The recombination activating proteins, RAG1 and RAG2 (RAG1/2), facilitate recombination by recognizing recombination signal sequences (RSSs) that flank the coding segments (4). Each RSS contains a relatively conserved heptamer and nonamer sequence separated by a poorly conserved 12 or 23 base-pair (bp) spacer sequence, known as the 12-RSS and 23-RSS, respectively. To stimulate recombination, a paired complex (PC) is formed consisting of a RAG1/2 heterotetramer bound to both a 12-RSS and a 23-RSS, in accordance with the “12/23 rule” (5,6). Upon PC formation, RAG1/2 cleaves DNA at RSS/coding junctions in a nick and hairpin-forming transesterification reaction, resulting in hairpin-sealed coding ends and open double-strand breaks at the 5’ ends of each RSS, termed signal ends (7). RAG1/2 releases the coding ends to nonhomologous end joining (NHEJ) factors and remains stably bound to the signal ends in a signal end complex (SEC) (8–10). The hairpin coding ends are opened and processed by addition or deletion of bases followed by coding end joining, which produces the exons encoding the variable region of the AgR. Signal ends are later joined heptamer-to-heptamer, typically with a precise junction, to form a signal joint (SJ).

In addition to shaping the AgR repertoire, V(D)J recombinase specificity has important implications for genomic stability. Off-target RAG-mediated cleavage events involving cryptic RSSs occur in a large subset of RAG-expressing pre-B-cells (11). Subsequent aberrant joining of the cleaved cryptic RSSs can result in chromosomal abnormalities and oncogenesis (12,13). Numerous co-factors can promote or inhibit V(D)J recombination, but recent studies have highlighted the importance of RAG1/2 DNA sequence specificity (14,15). A consensus RSS, CACAGTG-N_12/23_-ACAAAAACC, was derived from conservation of annotated RSSs (16). However, many RSSs in AgR loci diverge from the consensus RSS; therefore, RAG1/2 must be promiscuous enough to cleave divergent RSSs and generate AgR diversity, but it must also be precise to avoid cleaving cryptic RSSs and cause genomic instability.

We developed the **S**elective **A**mplification of **R**ecombination **P**roducts and **seq**uencing (SARP-seq) method to evaluate DNA selectivity in the V(D)J recombination reaction in an unbiased and high throughput approach. From the SARP-seq results described here, we provide the most comprehensive characterization of V(D)J recombinase specificity for RSSs to date. In a massively parallel format, these findings demonstrate RAG1/2 preference for selected purine/pyrimidine motifs in the RSS heptamer, elucidate effects of spacer sequence on recombination activity, and reveal DNA sequence-specific features that may enhance RAG-mediated cleavage through molecular dynamics simulations. Further, we demonstrate how SARP-seq results can be used to elucidate the relative contribution of RSS sequence to gene utilization at an endogenous AgR locus.

## RESULTS

### Selective Amplification of Recombination Products and Sequencing (SARP-seq)

The SARP-seq method (**Fig. 1A**) builds on a well-established approach that assays V(D)J recombination activity in RAG1/2-expressing cells using extrachromosomal plasmid substrates (17,18). Here, we coupled the extrachromosomal recombination assay with next generation sequencing (NGS) to simultaneously determine the relative utilization of thousands of RSSs in a complete V(D)J recombination reaction (**Fig. *1*B-E**). This was accomplished by randomizing the DNA sequence within an RSS to yield a large library of plasmid substrates (19). The utilization of each RSS in the recombination reaction was subsequently determined by NGS and a custom data analysis pipeline (Fig. S1).

**Figure 1.**
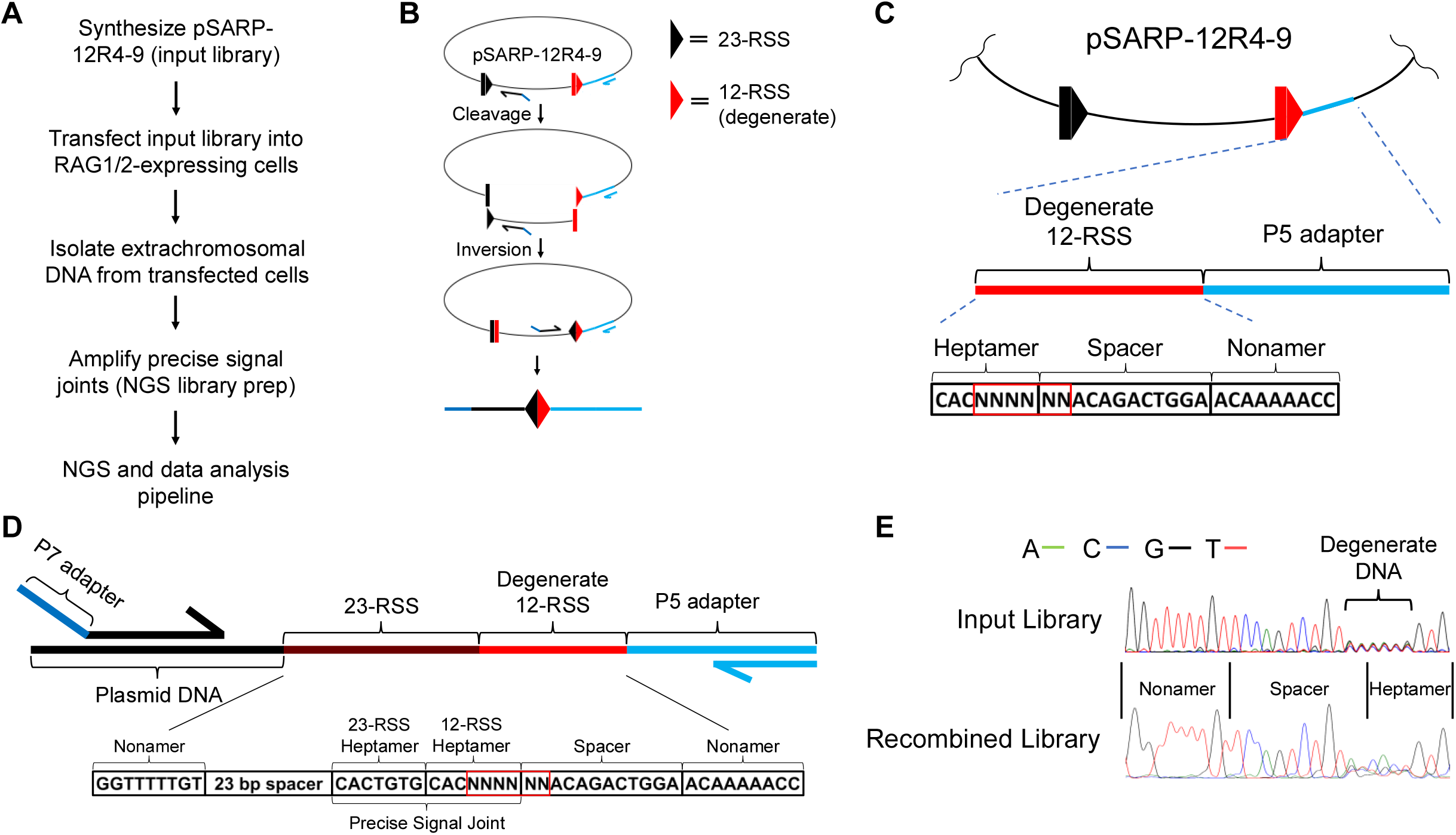
Schematic overview of SARP-seq method. **(A)** Flow chart of SARP-seq protocol **(B)** Diagram of episomal V(D)J recombination. The 12- and 23-RSSs are represented as red and black triangles, respectively. Half arrows represent PCR primers used to selectively amplify recombination products. The P5 adapter (light blue) and P7 adapter (dark blue) sequences are listed in *Table S3*. Coupled cleavage of 12- and 23-RSSs is followed by coding end (rectangles) joining and signal end joining by NHEJ. Joining of coding and signal ends results in an inversional recombination event. **(C)** Diagram of partially degenerate 12-RSS flanked by P5 adapter sequence. The non-degenerate spacer sequence is from murine VκL8 12-RSS, shown previously to impart good recombination efficiency (24). **(D)** Diagram of final PCR amplicon subjected to NGS. **(E)** Electropherogram charts of 12-RSS in pSARP-12R4-9 input library and of V(D)J recombination products (recombined library), showing the reverse complement sequence of the 12-RSS.

The plasmid substrate input library was constructed from the pMX-INV plasmid (17), which contains a 12-RSS and 23-RSS pair oriented in the same direction, with each RSS containing the consensus heptamer (CACAGTG) and nonamer (ACAAAAACC) sequences. To generate the plasmid substrate input library, a portion of the canonical 12-RSS was replaced with a fully degenerate sequence through directional ligation as described in *Materials and Methods* (**Fig. 1*C-D***). Illumina NGS adapter and sequencing primer sites were incorporated immediately upstream of the 12-RSS in pMX-INV to form an NGS-optimized plasmid library. Here, we focused on the heptamer and flanking spacer region of the 12-RSS (**Fig. 1*C***). The first three positions of the heptamer remained constant, as the CAC sequence is required for RAG-mediated DNA cleavage (20). The next six consecutive base-pairs, including 4 bp at the 3’-end of the heptamer plus the first 2 bp of the spacer, were fully randomized with equal ratios of A:C:G:T at each position. These positions correspond to bases 4-9 of the 12-RSS. Full degeneracy of positions 4-9 in the resulting plasmid input library, referred to as the pSARP-12R4-9, was confirmed by Sanger DNA sequencing (**Fig. 1*E***). The pSARP-12R4-9 library contains all 256 possible CAC-containing heptamer sequences, with each heptamer in turn flanked by 16 different adjacent spacer sequences. Altogether, the expected library size is a total of 4096 plasmid sequences; however, due to experimental constraints detailed in *SI Methods*, 4094 total sequences are expected.

V(D)J recombination of the pSARP-12R4-9 library leads to inversional rearrangement with the resulting signal and coding joints retained in the plasmid (**Fig. 1*B***). To initiate V(D)J recombination activity, the pSARP-12R4-9 library and core RAG1 and RAG2 expression vectors were co-transfected into 293T cells. Plasmid DNA was isolated 72 hrs after co-transfection, allowing sufficient time for extrachromosomal V(D)J recombination (Fig. S1*A*). The recombined output library was generated by PCR amplifying RSS signal joints(**Fig. *1B, D* and *E***). Sanger sequencing showed partial loss of degeneracy of the 12-RSS consistent with RAG1/2 selection of certain RSSs in V(D)J recombination (**Fig. *1E***). The output library was then subjected to NGS, and read counts for RSSs that formed precise signal joints were tabulated.

SARP-seq experiments using the pSARP-12R4-9 input library were replicated 3 times, and the resulting libraries were sequenced twice on the iSeq 100 platform and once on the miSeq platform. We refer to our first and second iSeq runs as “iSeq 1” and “iSeq 2”, respectively, and the miSeq run is referred to as “miSeq 1”. For iSeq 1 and iSeq 2, the entire SARP-seq protocol was repeated starting with preparation of the pSARP-12R4-9 input library, and miSeq 1 utilized the same degenerate 12-RSS input library as iSeq 2. The SARP-seq output libraries were sequenced at sequencing depths varying by >2 orders of magnitude. 12-RSSs utilized during V(D)J recombination were analyzed using a custom data analysis pipeline (Fig. *S1E* and described in *Materials and Methods*). In brief, reads containing a precise signal joint were extracted from FASTQ files, as precise signal joints are a hallmark of V(D)J recombination (**Fig. 1*D***). Differential read counts of precisely joined 12- and 23-RSSs correspond to V(D)J recombination frequencies. 12-RSS logos generated from precise signal joint read counts show striking reproducibility (**Fig. 2*B***). Information content for each position of the RSS heptamer was highly reproducible, and read counts for specific RSS motifs were reproducible across each independent SARP-seq experiment (**Fig. *2A* and *B***).

**Figure 2.**
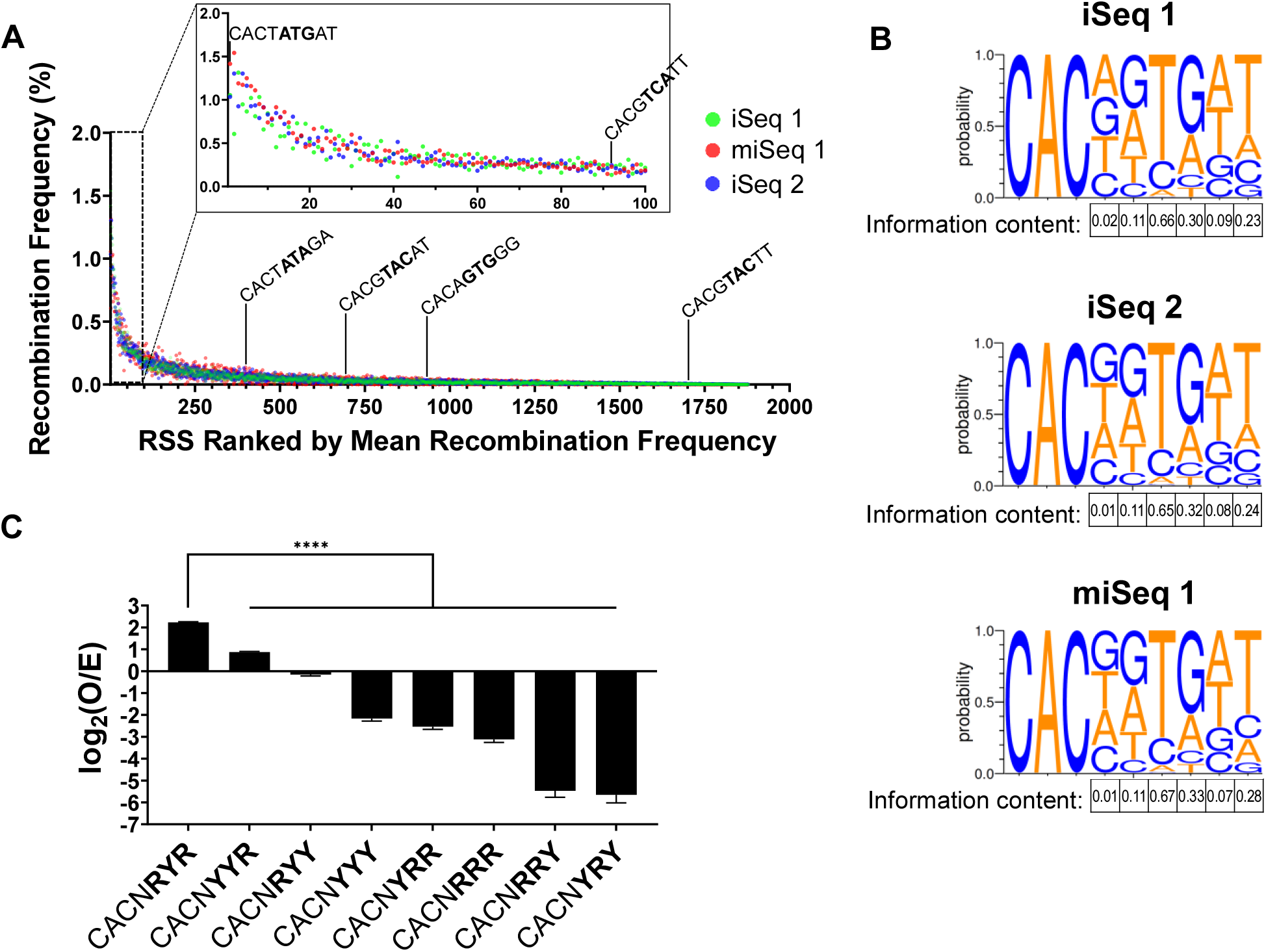
DNA sequence selectivity determined by SARP-seq. SARP-seq was replicated 3 times on two different sequencing platforms: twice on the iSeq 100 (iSeq 1 and iSeq 2) and once on the miSeq (miSeq 1). **(A)** Recombination frequencies of individual RSSs expressed as a percentage of total recombination events. Every RSS with reproducible V(D)J recombination activity was ranked by mean recombination frequency, so more efficacious RSSs occupied higher ranks and less efficacious RSSs occupied lower ranks. Efficacy was determined by calculating the mean recombination frequency of all three replicates (*Supplementary Table S1*). Specific RSSs are indicated by black pointers and listed from most efficacious to least efficacious: The top-ranked consensus R/Y motif (CACTATGAT), top-ranked RSS that completely lacks canonical consensus RSS base identity for heptamer positions 4-7 (CACGTCATT), median-ranked consensus R/Y motif (CACTATAGA, top-ranked anti-consensus R/Y motif (CACGTACAT), bottom-ranked canonical consensus RSS (CACAGTGGG), and median ranked anti-consensus R/Y motif (CACGTACTT). The top 100 RSSs are magnified in the panel inset. **(B)** Sequence logos depicting RAG1/2 specificity expressed as the probability of finding each base in precise signal joints. Total information content for each position of the degenerate 12-RSS region was calculated and expressed in nats. **(C)** Bar chart showing V(D)J recombination frequency of different purine/pyrimidine sequence motifs. Recombination frequencies are expressed as log_2_(O/E). *O* is the observed frequency of recombination events, and *E* is the expected frequency of random, non-specific recombination. Positive values indicate positive selection, and negative values indicate negative selection. Statistically significant differences were determined by ordinary one-way ANOVA with Dunnett’s multiple comparisons test (n = 3) (****, *p* < 0.0001).

Analysis of the SARP-seq results revealed a clear hierarchy of 12-RSSs utilized in V(D)J recombination (**Fig. 2 *A* and *C***, and Fig. S2). For example, >1800 unique 12-RSSs were found to reproducibly form a precise signal joint with the canonical 23-RSS, and the top 20 sequences made up 16.7% of all precise signal joints (Dataset S1). Conversely, more than 1600 unique sequences were poorer substrates that each made up less than 0.1% of total precise signal joints. Those sequences accounted for 40.1% of all precise signal joints despite accounting for 87.4% of unique sequences that reproducibly form precise signal joints (Dataset S1).

### The R/Y pattern in the RSS heptamer is highly predictive of RAG1/2 activity

By including all CAC-containing heptamers, SARP-seq deciphered heptamer features that increase V(D)J recombination. Five unique heptamer sequences encompassed 5.7% of total reads for precise signal joints (Supplementary Dataset S1). Four of these sequences differed only in position 4 of the CACNGTG motif. Interestingly, in addition to CACNGTG, the CACTATG heptamer was consistently in the top 2-3 heptamers used (**Fig. 2*A*** and Supplementary Dataset S1). CACTATG is the heptamer sequence in the terminal inverted repeat (TIR) of the ProtoRag transposon identified from *Branchiostoma belcheri* (Bb) (21). While mammalian RAG enzymes efficiently cleave RSS substrates containing the Bb RSS heptamer, this heptamer sequence is poorly represented in endogenous RSSs at AgR loci in both mice and human (22).

The sequence logos from each experiment represent the base preference at each position (**Fig. 2*B***). Based on the sequence logo, position 4 in the RSS heptamer showed the least discrimination of base identity amongst all the heptamer base positions (**Fig. 2*B***). The preference for heptamer positions 5-7 largely depended on the purine (R) or pyrimidine (Y) classification of each base (**Fig. *2 B* and *C*)**. For example, R (A or G) was preferred over Y (C or T) at positions 5 and 7 by approximately 2-fold and 7-fold, respectively (**Fig. 2 *B* and *C***). Notably, the R/Y specificity was most pronounced at position 6, where RAG1/2 preferred Y over R by greater than one order of magnitude (**Fig. 2 *B* and *C*)**. Consistent with a previous biochemical study, G is least preferred at position 6 (23), and >80% of RSS heptamers containing G in this position are in the bottom fourth of RSSs reproducibly utilized in the SARP-seq method (**Fig. 2*B*** and Supplementary Dataset S1).

Normalized recombination frequencies for every RSS studied were ranked by their mean recombination frequency with the more efficacious RSSs ranked higher than less efficacious RSSs (**Fig. 2*A***). The top-ranked and median-ranked consensus R/Y motifs (CACNRYR) were ranked #1 and #400, respectively. In contrast, the top-ranked and median-ranked anti-consensus R/Y motifs (CACNYRY) were ranked #694 and #1652, respectively. While the consensus R/Y motif is a feature shared with the canonical consensus RSS (CACAGTG), the consensus R/Y motif provides a much more accurate, comprehensive description of V(D)J recombination specificity. The canonical consensus RSS describes less than 5% of V(D)J recombination products, and the canonical consensus heptamer is only the 933^rd^ most efficacious RSS when flanked by a “GG” spacer sequence (**Fig. 2*A*** and Dataset S1). Interestingly, the sequence “CACGTCATT” shares no base identity with canonical consensus RSS at heptamer positions H4-H7 but is a top-100 12-RSS, vastly outcompeting the canonical consensus RSS sequence “CACAGTGGG” (**Fig. 2*A***). The consensus R/Y sequence, however, describes more than 50% of all V(D)J recombination products (**Fig. 2*C***). We quantified RSS utilization for every R/Y motif spanning positions 5-7 (**Fig. 2*C***). R/Y specificity is striking, with RAG1/2 preferring CACNRYR > 100-fold more than CACNYRY (**Fig. 2*C***). Nevertheless, the “consensus” R/Y motif CACNRYR was only enriched by ~4-fold relative to simulated random RSS selection (**Fig. 2*C*)**. Most R/Y motifs were depleted by >4-fold compared to random sequence selectivity, with >16-fold depletion for CACNRRY and CACNYRY (**Fig. 2*C***). Thus, while RAG1/2 does not strongly select for a specific consensus motif, it strongly selects against some R/Y motifs (**Fig. 2*C***). We also quantified RAG1/2 activity for every possible combination of “weak” A/T (W) and “strong” G/C (S) sequences (Fig. S2*D*). RAG1/2 favored CACNSWS by < 10-fold over the least active S/W motif, CACNSSW (Fig. S2*D*), demonstrating that the R/Y pattern is a more accurate representation of RAG1/2’s sequence motif preference. To further validate RAG1/2 preference for CACNRYR over CACNYRY, we performed low-throughput extrachromosomal V(D)J recombination assays by cloning an active consensus (CACAGTGAT) and anti-consensus (CACGTACAT) RSS into pMX-INV and performing semi-quantitative V(D)J recombination assays (*Materials and Methods* and Fig. S2*E*). A statistically significant difference in RAG1/2 activity was observed for the same consensus and anti-consensus RSSs in both the low-throughput assay and SARP-seq (Fig. S2*E*).

### The heptamer-flanking spacer sequence is a major determinant for RSS utilization

While the 12 and 23 bp spacers are the least conserved regions of the RSS, previous studies showed some DNA sequence preferences at the 5’ end of the spacer in the 12-RSS (24,25). Thus, we randomized the first two positions of the 12 bp spacer in the SARP-seq pSARP-12R4-9 input library, such that each RSS heptamer is sampled with 16 different spacer sequences. The SARP-seq results can be summarized in two main points. First, the relative ranking of the 16 spacers show that the top 4 ranking spacers all include T at the 2^nd^ position (position 9 in the 12-RSS), and a preference for A or T at the first position of the spacer (**Table 1** and Fig. S3 *A* and *B*). The poorest spacers are G/C rich, with CG and GG as the bottom two ranked spacer sequences (**Table 1** and Fig. S3 *A* and *B*). Second, the adjoining spacer sequence has profound effects on heptamer utilization (**Table 1** and Fig. S3*C-F*). For example, the frequency at which the consensus CACAGTG heptamer is utilized varies considerably depending on the adjoining spacer sequence, with a >50-fold difference in observed/expected values for the CACAGTG consensus heptamer flanked by a ‘good’ versus a ‘poor’ spacer (**Fig. 2*A*** and Dataset S1). The frequency with a ‘poor’ spacer brings the utilization of the CACAGTG heptamer to similarly low levels as sequences containing anti-consensus CACNYRY or CACNRRY heptamer motifs (**Fig. 2*A***).

**Table 1.** Top RSS heptamer sequences for each spacer sequence falling in the top quartile (upper tables) or bottom quartile (lower tables) of recombination frequency

| AT Spacer |  | TT Spacer |  | GT Spacer |  | CT Spacer |  |
| --- | --- | --- | --- | --- | --- | --- | --- |
| ** CACTATGAT | 1.168 | CACTGTGTT | 1.141 | CACTATGGT | 0.516 | CACAGTGCT | 1.150 |
| * CACAGTGAT | 1.144 | CACGGTGTT | 0.925 | CACAGTGGT | 0.452 | CACTATGCT | 0.557 |
| CACTGTGAT | 1.093 | CACAGTGTT | 0.829 | CACTGTGGT | 0.448 | CACTGTGCT | 0.515 |
| CACGGTGAT | 1.036 | CACTATGTT | 0.805 | CACGGTGGT | 0.344 | CACGGTGCT | 0.499 |
| CACCGTGAT | 0.952 | CACCGTGTT | 0.721 | CACAATGGT | 0.344 | CACCGTGCT | 0.442 |
| CACCATGAT | 0.755 | CACGTTGTT | 0.620 | CACGATGGT | 0.338 | CACAATGCT | 0.330 |
| CACGATGAT | 0.739 | CACAATGTT | 0.583 | CACCATGGT | 0.328 | CACCATGCT | 0.317 |
| CACAATGAT | 0.726 | CACGATGTT | 0.516 | CACCGTGGT | 0.313 | CACGTTGCT | 0.247 |
| CACGTTGAT | 0.680 | CACATTGTT | 0.508 | CACGTTGGT | 0.243 | CACGATGCT | 0.228 |
| CACATTGAT | 0.597 | CACCATGTT | 0.504 | CACTGTAGT | 0.242 | CACTATACT | 0.157 |
| GG Spacer |  | CG Spacer |  | CC Spacer |  | GA Spacer |  |
| CACGTTGGG | 0.066 | CACTGTGCG | 0.103 | CACCGTGCC | 0.131 | CACTATGGA | 0.106 |
| CACTATGGG | 0.064 | CACGGTGCG | 0.088 | CACAGTGCC | 0.128 | CACTGTGGA | 0.104 |
| CACGATGGG | 0.054 | CACCGTGCG | 0.085 | CACTGTGCC | 0.128 | CACGGTGGA | 0.084 |
| CACCGTGGG | 0.040 | CACGATGCG | 0.073 | CACTATGCC | 0.119 | CACGGTAGA | 0.075 |
| CACTGTGGG | 0.037 | CACTATGCG | 0.072 | CACGATGCC | 0.109 | CACCGTGGA | 0.066 |
| CACCATGGG | 0.031 | CACAGTGCG | 0.052 | CACGGTGCC | 0.090 | CACTATAGA | 0.062 |
| CACAATGGG | 0.029 | CACATTGCG | 0.041 | CACAATGCC | 0.089 | CACAGTGGA | 0.060 |
| CACGGTAGG | 0.027 | CACCATGCG | 0.040 | CACCATGCC | 0.072 | CACCATGGA | 0.054 |
| CACGGTGGG | 0.022 | CACAATGCG | 0.036 | CACGTTGCC | 0.050 | CACGTTGGA | 0.053 |
| CACAGTCGG | 0.021 | CACGTTGCG | 0.033 | CACATTGCC | 0.049 | CACAGTAGA | 0.051 |
| CACAGTGGG | 0.020 |  |  |  |  |  |  |

\* Canonical RSS heptamer sequences are highlighted.
\*\* Count frequency data from Supplementary Dataset S2.

### RSS R/Y motif adopts dynamic structural features unique to YpR base-pair steps

YpR steps can have unique base-pair step features that cause local distortions in a B-DNA helix (26,27). Recent cryo-EM and X-ray crystallographic data show the RSS heptamer untwists in the RAG1/2 active site prior to RSS cleavage (28–30). RAG-mediated cleavage therefore requires DNA flexibility, and RAG1/2 makes few base-specific heptamer contacts outside of the highly conserved CAC and dT at heptamer position 6 (29). While strong base-specific contacts with an intact, B-form RSS heptamer could enhance binding affinity and specificity, they could also hinder RAG1/2 catalysis by creating an energetic barrier to DNA structural transitions in the RAG1/2 active site. We hypothesized that the RSS heptamer motifs favored by RAG1/2 are structurally flexible, and RAG1/2 preference for R/Y motifs could be explained by dynamic structural features unique to RpY and YpR dinucleotide steps that accommodate heptamer untwisting in the RAG1/2 active site. To test this hypothesis, we performed *in silico* simulations of a highly active consensus R/Y RSS (CACAATGAT) and a poorly active anti-consensus R/Y RSS (CACTTATGT) (**Fig. 3**).

**Figure 3.**
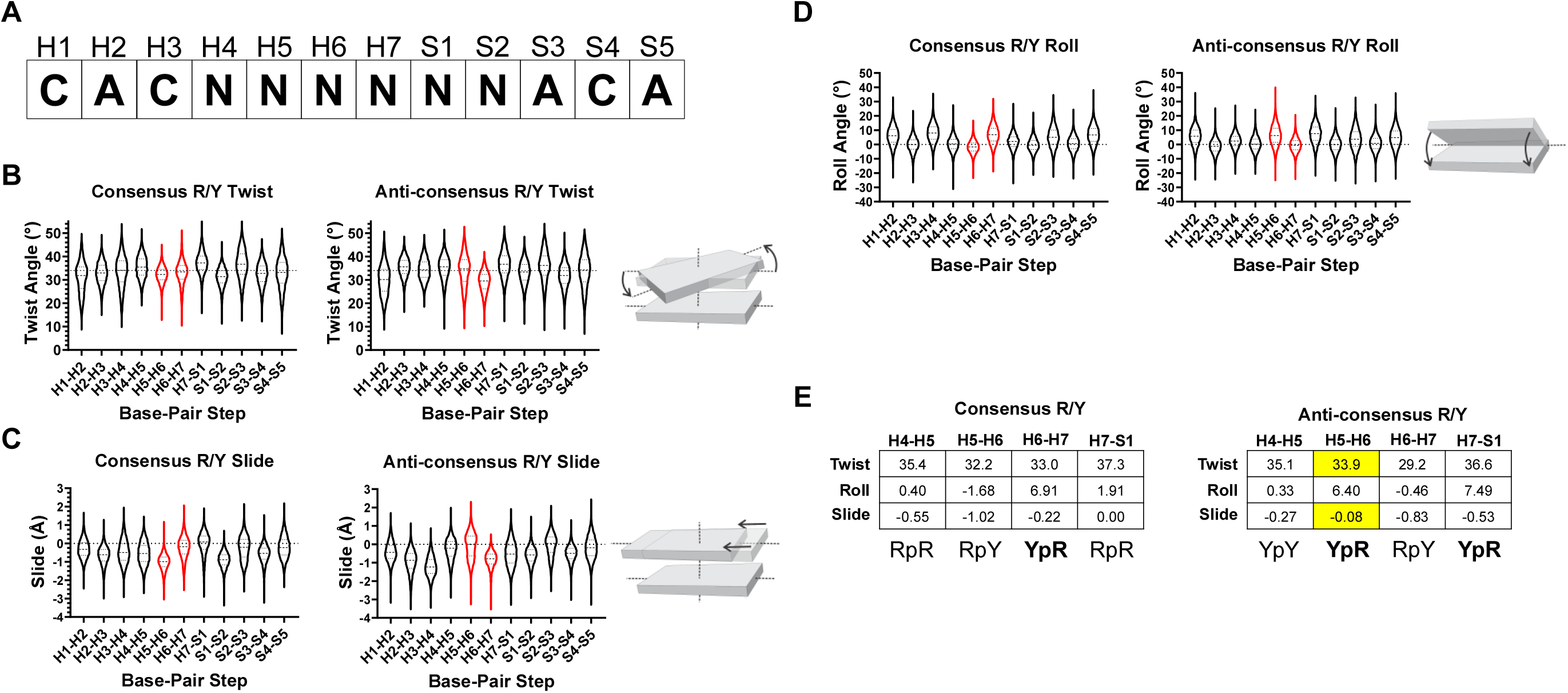
Molecular dynamics simulations of a consensus R/Y RSS (CACAATGAT) and an anti-consensus R/Y RSS (CACTTATGT). **(A)** Base position nomenclature for the 12-RSS heptamer and 5’ spacer region as used in subsequent panels. Violin plots depict probability distributions for **(B)** twist, **(C)** slide, and **(D)** roll. Base-pair steps between heptamer positions 5-7 are colored red. Diagrams illustrating each base-pair step parameter is shown adjacent to the corresponding plots. **(E)** Tables of mean twist, roll, and slide values for base-pair steps contiguous to heptamer positions 5-7. The R/Y composition for each base-pair step is shown below each row of the columns, with YpR in boldface. Asymmetric probability distributions are highlighted yellow.

The R/Y consensus adopted unique twist distributions compared to the anti-consensus and inactive RSSs (**Fig. 3*B*** and Fig. S4*C*). The first CpA base-pair step adopted twist-angle probability distributions that were asymmetric, a feature previously characterized in YpR steps (**Fig. 3 *B* and *E***, Fig S4*C*) (27). Most other YpR steps in simulated RSSs showed asymmetric twist-angle distributions characterized by an abundance of highly untwisted populations (**Fig. 3 *B* and *E***). An exception to this was the TpG step in the simulated R/Y consensus RSS heptamer, which did not have an asymmetric twist-angle distribution (**Fig. 3 *B* and *E***). We also measured base-pair slide and roll angle distributions for RSS base-pair steps (**Fig. *3C-E***). Consensus and anti-consensus RSS heptamers have markedly different slide and roll angle distributions for base-pair steps at, and proximal to, heptamer positions 5-7. The starkest contrast in roll and slide distributions are at the YpR/RpY steps at heptamer positions 5-6 and 6-7 (**Fig. 3*E***). In both consensus and anti-consensus RSSs, the YpR step has a large roll angle and little-to-no slide - features that can induce DNA bending and major groove compression (31). The YpR step in the anti-consensus RSS is offset by one base position, which may incorrectly orient DNA structural distortions, such as unwinding, needed for RAG-mediated cleavage (29,30). Because RAG1/2 makes minor groove contacts proximal to H4-6, we also measured RSS minor groove widths (Fig. S4 *A* and *B*).The consensus RSS heptamer had a wider range of minor groove widths, demonstrating its flexibility (Fig. S4 *A* and *B*). Minor groove flexibility may accommodate RAG1 interactions with the minor groove as well as helical distortions needed for heptamer unwinding (Supplementary Movie S1) (29,30). Minor groove widths in the anti-consensus RSS heptamer were more tightly distributed about smaller mean values, indicating a more rigid and narrow minor groove, particularly at H2-4 (Fig. S4 *A* and *B*). Because heptamer unwinding during RAG-mediated cleavage is caused by a combination of untwisting and helical distortions, we also measured minor groove widths when the first base-pair step (CpA) of the RSS heptamer is untwisted (H1-H2 twist angle <17°) to characterize twist-coupled minor groove flexibility. Interestingly, the anti-consensus minor groove appears even more rigid and narrow when H1-H2 is untwisted, whereas the consensus RSS heptamer minor groove remains flexible. This suggests that the consensus R/Y motif better supports groove and base-pair step parameter flexibility than the anti-consensus R/Y motif, including twist-coupled minor groove deformations.

### Comparative analysis of SARP-seq results to RAG1/2 activity on endogenous RSSs

Statistical models for RSS recombinogenic potential were previously developed (32,33), providing a scoring system for RSS information content (RIC). We calculated RIC scores for all RSSs observed in SARP-seq to compare RIC score to RSS efficacy determined by SARP-seq (**Fig. 4 *A* and *B***). RIC scores were positively correlated with RSS efficacy (Spearman r = 0.47) (**Fig. 4 *A* and *B***). Despite this, RIC was only somewhat predictive of RSS efficacy (**Fig. 4 *A* and *B***). RIC scores were derived from conservation of endogenous RSSs, as opposed to relative RAG activity at each RSS. This likely accounts for the relatively poor correlation between RIC scores and SARP-seq results (**Fig. 4 *A* and *B***), indicating that the importance of positions 4-9 in the 12-RSS for determining probability of RAG-mediated cleavage has been vastly underestimated.

**Figure 4.**
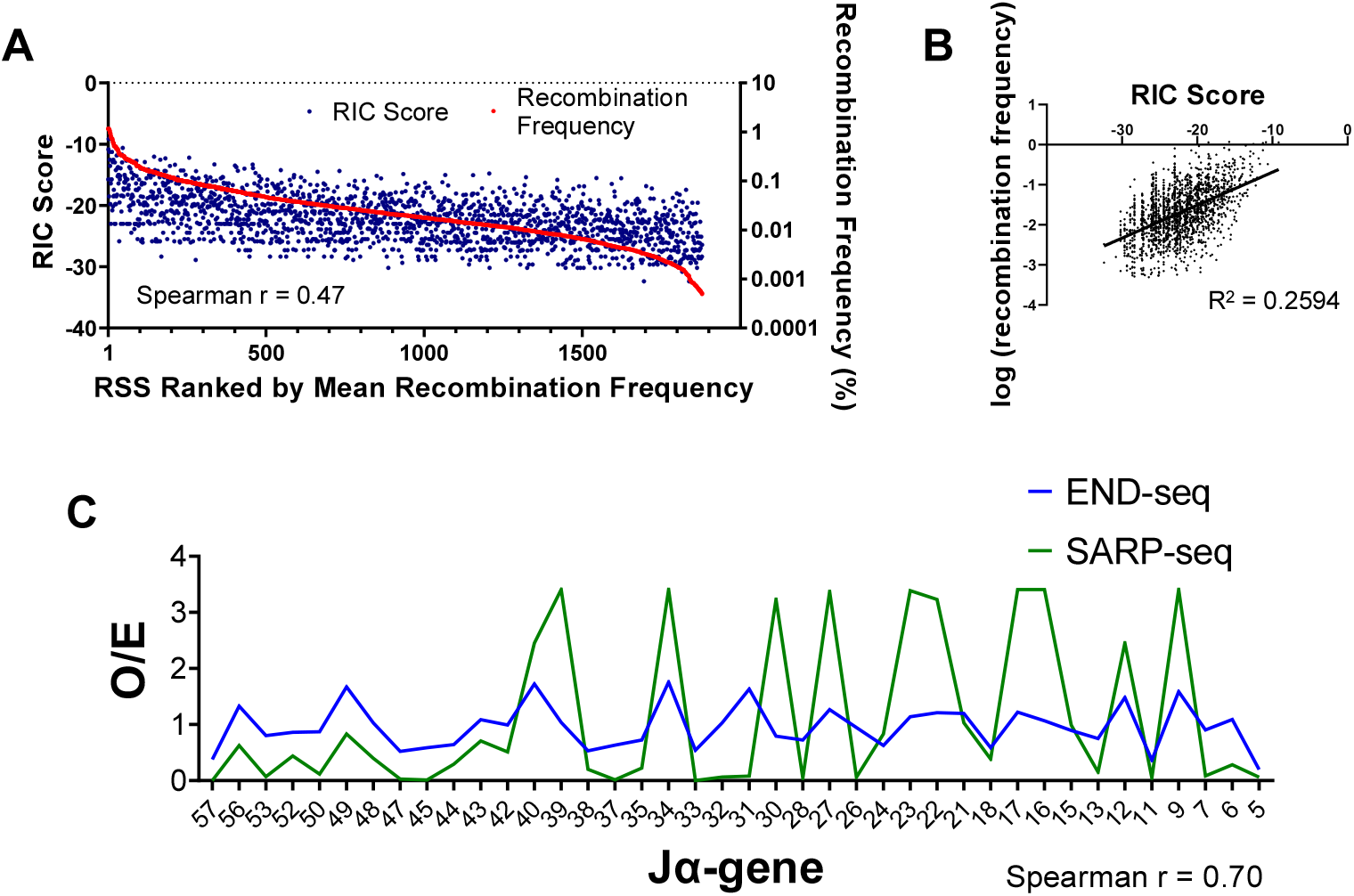
Endogenous RSS information content (RIC) scores and endogenous *Tcra* J-gene recombination compared with SARP-seq recombination. **(A)** Calculated RIC scores (blue dots, left y-axis) for each RSS analyzed in SARP-seq. Red line indicates mean RSS recombination frequency for each RSS characterized by SARP-seq (right y-axis, n = 3), and RSSs were ranked along the x-axis with most efficacious RSSs occupying higher ranks and less efficacious RSSs occupying lower ranks. **(B)** Log-transformed mean recombination frequencies for each RSS characterized by SARP-seq (n = 3) were plotted against RIC score. Black trend-line was generated using a linear regression model of log-transformed RSS recombination frequency expressed as a function of RIC score. **(C)** Mean normalized SARP-seq count frequencies and normalized END-seq count frequencies expressed as *O/E* where *O* is the observed frequency and *E* is the expected frequency if V(D)J recombination is completely random and nonspecific. END-seq expected frequency *E* was calculated by dividing the total END-seq counts for each RSS included in the analysis by the total number of unique RSSs being counted. Chromosomal position-specific effects were accounted for in the END-seq data by analyzing RAG cleavage efficiency of each J-gene relative to the cleavage efficiency of two flanking J-genes on either side (quantifications are provided in Supplementary Dataset S2).

RSS quality has been challenging to predict based on DNA sequence, since endogenous RSSs are embedded in a range of chromatin environments that can affect selectivity of individual RSSs by RAG1/2 (34,35). Examples of sequence-independent effects on RSS utilization include nucleosome density and positioning relative to the RSSs (36); the presence of certain histone marks, such as H3K4me3, that serve as a chromatin docking site for full length RAG2 (37); and 3-dimensional chromatin organization that can impact RAG1/2 access to RSSs (38). Here, we tested how the SARP-seq results correlated with RSS cleavage at an endogenous AgR locus. Specifically, we compared our SARP-seq results to the relative frequency of RAG-mediated cleavage across the murine *Tcra* J_α_ region, which was previously reported using the END-seq method (11). In this 64 kb region, there are 48 12-RSSs with CAC-containing heptamers (**Fig. *4C*** and Supplementary Dataset S2). Further, the *Tcra* J_α_-associated RSSs show considerable sequence variability through the heptamer and adjoining spacer region, providing an excellent test set to compare against the SARP-seq results. Previous studies have shown that the *Tcra* V_α_-to-J_α_ rearrangement is highly processive with attempts to join proximal V_α_ to the most 5’ J_α_ segments occurring prior to rearrangement of more distal 3’ J_α_ gene segments (39,40). These findings suggest RSS quality, based on DNA sequence, would not significantly factor into gene segment utilization. Nevertheless, we determined that there is a statistically significant correlation (Spearman correlation = 0.70, *p* < 0.0001) between RSS utilization for the END-seq versus SARP-seq results (**Fig. 4*C*** and Supplementary Dataset S2). This is striking, considering that only the heptamer and the first 2 bp of the spacer can be exactly matched. RSS quality was generally better for more distal RSSs and poorer for more proximal RSSs (**Fig. 4*C*** and Supplementary Dataset S2). The proximal RSSs would be within or near RAG-enriched recombination centers (39,40). Despite higher quality RSSs, more distal 12/23-RSS pairs were less frequently utilized during V(D)J recombination *in vivo* (39,40). These data suggest sequence-based RSS quality can help compensate for less frequent encounters of distal 12/23-RSS pairs. In summary, by comparing SARP-seq results from extrachromosomal substrates lacking specific chromatin modifications to chromosomal V(D)J recombination sequence specificity *in vivo*, it may be possible to discern *in vivo* RAG1/2 targets driven primarily by DNA sequence specificity from targets that are recombined due to epigenetic RAG1/2 effectors.

## DISCUSSION

Our development and implementation of SARP-seq, a high throughput V(D)J recombination assay, revealed RAG1/2 DNA sequence preference for every functional 12-RSS heptamer sequence containing a “CAC”. These results provided a hierarchical ranking of RSS heptamers targeted by RAG1/2, showing that a broad range of heptamer sequences can be utilized in V(D)J recombination. The effect of the first two positions of the 12-RSS spacer on RAG1/2 activity was also assayed, demonstrating the range of recombination frequencies for each RSS heptamer that occurred with 16 separate flanking spacer sequences. Previous studies showed RAG1/2 can cleave non-consensus RSSs (11,41–45), and the abundance of genomic non-consensus RSSs necessitates promiscuous RAG1/2 activity to include every functional antigen receptor gene in the antigen receptor repertoire.

RAG1/2 showed a striking lack of specificity for adenine at heptamer position 4. This lack of specificity is consistent with recent structures of RAG:RSS complexes that showed little to no contact between RAG1/2 and heptamer position 4 (29). While it is possible that the non-core regions of RAG1/2, which are absent in the constructs used here, could affect preference at heptamer position 4, no previous studies have shown a role for the non-core regions in sequence-specific DNA interactions. Further, cryo-EM and X-ray crystallographic studies that included non-core RAG1/2 domains showed no resolvable cryo-EM/electron density for non-core RAG1/2 but extensive RAG-DNA contacts between the RSS and core RAG1/2 domains (28–30,46). RAG1/2 also showed weak A/T preference at the first two positions of the RSS spacer, implying a nonamer rather than heptamer RAG1/2 recognition sequence. Interestingly, extending the consensus RSS through the first two positions of the putative spacer yields a nonamer-nonamer recognition site separated by 10 bp, which corresponds precisely to one turn of B-form DNA helix (at 10.5 bp) for the spacer DNA rather than 12 bp for the current heptamer-nonamer classification of the 12-RSS. Future SARP-seq experiments extending further into the RSS spacer are needed to validate the nonamer-spacer-nonamer RSS model.

RAG1/2 targeted a surprisingly large variety of sequences spanning heptamer positons 5-7. Nevertheless, there was a clear preference for certain purine/pyrimidine (R/Y) motifs. R/Y RSS motifs containing a YpR step at heptamer positions 6-7 were the only enriched R/Y motifs found in precise signal joints (**Fig. 2*C***), highlighting its importance for RAG1/2 cleavage compared to other RSS heptamer features. Notably, YpR dinucleotides have low base-stacking energies and are among the most conformationally flexible base-pair steps (27,47,48). The deformability of YpR steps allows them to act as a flexible “hinge” during protein:DNA interactions (48–50). Prior to nicking, RAG1/2 contacts T at heptamer position 6 through a wide minor groove, which explains the preference for T over C. The RSS heptamer must unwind in the RAG1/2 active site to correctly position the scissile phosphate next to the catalytic residues (28). While the unwinding is mostly localized to the first 3 bases of the RSS heptamer, heptamer positions 4-7 are under-twisted, and the minor groove remains widened (29,30). The RSS heptamer must be flexible enough to accommodate these structural distortions prior to nicking. As RAG1/2-mediated DNA cleavage is ATP-independent, the RSS heptamer must be able to unwind in the RAG1/2 active site without an external force driving the structural transition. These observations fit a model in which R/Y heptamer motif preference of RAG1/2 is a result of RAG1/2 selecting for DNA sequences that can accommodate the structural transitions necessary for RAG-mediated cleavage.

The quality of RSSs may have evolved to modulate RAG1/2 activity at certain locations within AgR loci. In addition to the poor quality 5’ J_α_ RSSs in the mammalian *Tcra* locus (**Fig. 4*C***), the poor quality of V_β_ RSSs was proposed to reinforce allelic exclusion of *Tcrb* by limiting RAG1/2 activity. In a previous study, replacing a poor quality V_β_ RSS with a consensus RSS led to a dramatic increase in utilization of the adjacent V_β_ gene, as well as increased allelic inclusion, demonstrating the effect of RSS quality on V(D)J recombination efficiency in the context of an AgR loci (14,15). Still, it has been challenging to discern the contribution of RSS quality to RAG1/2 activity in varying chromatin environments. RAG-RSS interactions in AgR loci are also proposed to be driven by RAG1/2 scanning through cohesin-mediated DNA loops. Regardless of whether RAG recognizes RSSs through linear DNA scanning or three-dimensional diffusion, RSS quality could enhance or repress RAG-mediated cleavage by accommodating or resisting heptamer unwinding prior to nicking and hairpin formation. The R/Y motif preference in the heptamer determined by the SARP-seq results, along with evidence of heptamer base unwinding in RAG-RSS high resolution structures, are consistent with the importance of this region of the RSS in affecting the rates for nicking and hairpin formation. However, we cannot rule out that a subset of RSSs is less efficiently utilized due to defects in NHEJ in forming precise signal joints, which may occur if RAGs release particular RSSs before proper handoff to NHEJ factors. Future studies will be needed to delineate RAG1/2 cleavage from NHEJ.

Our comprehensive characterization of V(D)J recombinase activity on RSS heptamer sequences revealed preference for conformationally flexible R/Y motifs and even stronger selection against RSS heptamers with an RpY step at heptamer positions 6-7. Given the considerable sequence divergence of RSSs at some AgR loci, RAG1/2 specificity by exclusion may facilitate full AgR repertoire realization without compromising genomic stability. The present SARP-seq study focused on V(D)J recombination specificity for positions 4-9 of the 12-RSS heptamer/spacer region. The flexibility of this method will facilitate future studies on a broad range of questions regarding DNA selectivity in V(D)J recombination.

## MATERIALS AND METHODS

### Input library preparation

To generate the SARP-seq input plasmid substrate library, pMX-INV (kindly provided by Barry Sleckman) was digested with MluI and EcoRI, and subsequently purified by extraction from a 1% agarose gel using the Monarch DNA Gel Extraction Kit following manufacturer instructions (NEB # T1020S). The partially degenerate DNA oligonucleotide, 12RSS4-9 (sequence in Supplementary Table S1), was duplexed with Duplexing Primer (sequence in Supplementary Table S1) by a single annealing and primer extension step with Q5 high-fidelity polymerase. Duplexed 12RSS4-9 was digested with MluI and EcoRI and purified using the Zymo DNA Clean and Concentrator Kit (Zymo # D4013). The double-digested and gel purified pMX-INV and duplexed 12RSS4-9 were ligated at a 3:1 12RSS4-9:pMX-INV ratio with T4 DNA Ligase by overnight incubation at 16°C, and purified using the Zymo DNA Clean and Concentrator kit. The resulting ligation product, pSARP-12R4-9, was used as an input library for extrachromosomal V(D)J recombination experiments in the SARP-seq protocol.

### Extrachromosomal V(D)J recombination experiment and SARP-seq

HEK293T cells were seeded in a 10 cm plate and grown overnight to 60-75% confluency in media containing DMEM, 10% FBS, 1x antibiotic-antimycotic, and 1 mM sodium pyruvate. The cells were then co-transfected with an expression vector for maltose binding protein core-RAG1 fusion protein (MBP-cRAG1) (kindly provided by Patrick Swanson), an expression vector for mCherry-tagged core-RAG2 (Ch-cRAG2) (51), and the pSARP-12R4-9 input library with a 3:1 transfection reagent:DNA ratio using Fugene 6 transfection reagent (Promega #E269A). At 72 hrs, ~90% of cells were transfected, as determined by quantifying the proportion of cells expressing mCherry. Plasmid DNA was then purified using a modification of the Hirt preparation (52). Briefly, the cells were harvested by trypsinization, washed in PBS, and lysed by resuspension in digestion buffer (10mM Tris pH 7.5, 100mM NaCl, 25mM EDTA, 0.5% SDS, 0.1mg/mL proteinase K) overnight at 37°C. Cell lysate was equilibrated to 1 M NaCl by addition of 5 M NaCl solution, incubated overnight at 4°C, and the supernatant retained following centrifugation. The plasmid DNA in the supernatant was subsequently concentrated by ethanol precipitation. Signal joints present in the recovered plasmid were selectively amplified with a nested PCR approach using Q5 High-Fidelity DNA Polymerase. Primer “Nest FWD” and “Nest RVS” were used and the PCR amplicon was PAGE purified. Next, primers “P7 primer” and “iSeq 1 P5 primer” were used to amplify the iSeq 1 library while primers “P7 primer” and “miSeq P5 primer 1” were used to amplify the miSeq 1 and iSeq 2 libraries. The PCR products for miSeq 1 and iSeq 2 libraries were PAGE purified and PCR amplified with primers “P7 primer” and “miSeq P5 primer 2.” Primer sequences are provided in Supplementary Table S1. The final PCR products for iSeq 1, iSeq 2, and miSeq 1 libraries were PAGE purified and subjected to Illumina next generation sequencing on the iSeq 100 or miSeq platform (iSeq runs were performed at the OUHSC Nathan Shock Center of Excellence in the Biology of Aging, and the miSeq run was performed at the OUHSC Laboratory of Molecular Biology and Cytometry Research).

### Analysis of next-generation sequencing data

FASTQ sequences with mean Q < 20 were filtered out with PRINSEQ (53). “grep” was used to match filtered reads containing a precise 12/23 signal joint sequence (regex:….GTGCACAGTG), and matched reads were written to a new FASTQ file. Reads containing mismatches in the 12-RSS nonamer were filtered out by matching the consensus nonamer sequence with “grep” and writing matching reads to a new file in FASTQ format. Next, all bases except the 12-RSS heptamer and the first 2 nucleotides of the 12-RSS spacer were trimmed using the “trim” tool implemented in the Galaxy web platform (54). Trimmed reads with mean Q < 30 were filtered out. Sequences were converted to FASTA format, reversed, and complemented. RSS heptamers without a “CAC” were filtered out by matching lines beginning with “CAC” and writing matched reads to new file in FASTA format. Unique occurrences of each RSS sequence were counted to form count tables for each sequencing run. Counts for each unique, recombined RSS were expressed as a percentage of total recombined RSS counts to account for differences in sequencing depth.

### Molecular dynamics simulations

Initial models for molecular dynamics simulations were generated by modeling recombination signal sequences into a B-DNA structure using the “fiber” command within X3DNAv2.4 (55). B-DNA was simulated using the parmbsc1 force-field implemented in GROMACS-2019 (56–58). The RSS models were placed in a solvent box containing a 10 nm buffered layer of explicit tip3p water molecules. The solvent box was neutralized by randomly replacing solvent water molecules with potassium ions. The system was subject to energy minimization using < 50,000 iterations of a steepest descent algorithm or until the system force < 1000 kJ/mol/nm. The minimized structure was used as an input for 100 ps temperature equilibration to 300 K using a weak-coupling modified Berendsen thermostat. The temperature equilibrated system was used as an input for 100 ps of pressure equilibration to 1 bar using an isotropic Parrinello-Rahman barostat. After pressure equilibration, the system was subject to 100 ns of production simulation time. Simulations were run with a leapfrog integrator over a 2 fs time step. Hydrogen bonds were constrained by 4^th^ order lincs algorithm. Nonbonded interactions were calculated using a Verlet cutoff scheme with a 1 nm cutoff for both short-range electrostatic interactions and short-range Van der Waals interactions. Particle Mesh Ewald summation with 0.15 nm grid spacing was used to calculate long-range electrostatic interactions.

### Analysis of molecular dynamics data

Post-production processing and analysis of molecular dynamics simulation data was performed using GROMACS-2019 and X3DNA software packages (55,56,58). Molecules of interest were centered in the solvent box, and trajectory files were converted to PDB format. The first 20 ns of simulation time were excluded from further analysis to allow for equilibration. A PDB file containing coordinates for each simulated time-step was used as an input for X3DNA’s “analyze” program, which generated a complete, reversible set of helical and base-pair step parameters capable of rebuilding the DNA molecules of interest (55). Base-pair step parameters and minor groove widths for each time-step were used to generate **Fig. 4** and Supplementary Fig. S4. Minor groove widths were calculated as a simple inter-phosphate distance using the X3DNA software package (55,59).

### PAGE purification

The PCR product was separated on a 10% polyacrylamide gel for ~1 h at 125 V with a cooling circulating water bath. The gel was stained with 1x SYBR safe (Thermofisher #S33102), visualized under blue light, and the band of interest was excised from the gel. The PCR product was eluted from the gel slice by crushing the gel slice and soaking in TE buffer (pH 7.5) buffer overnight (“crush-and-soak” method). The PCR product was further purified using a Zymo DNA Clean and Concentrator Kit (Zymo # D4013).

### Low throughput extrachromosomal V(D)J recombination assay

The 12-RSS in pMX-INV was replaced with a consensus (CACAGTG-AT) or anti-consensus (CACGTAC-AT) 12-RSS. The constructs were separately transfected into HEK 293T cells with the MBP-core-RAG1 and mCherry-core-RAG2 expression vectors, and following 72 hrs of cell growth, the plasmid DNA was purified as in the SARP-seq protocol. Purified plasmid DNA was serially diluted by 2-fold three times and subjected to PCR amplification of total pMX-INV DNA and V(D)J recombined DNA using primers “Nest FWD” and “Nest RVS” to amplify V(D)J recombined pMX-INV DNA and primers “Nest FWD” and “Input RVS” to amplify total input pMX-INV DNA (Supplementary Table S1). PCR products were separated on an 8% polyacrylamide gel, stained with SYBR safe, and imaged. Band intensities were quantified, and recombination activity was quantified by dividing recombined DNA band intensity by input DNA band intensity.

### Statistics

Statistical tests were performed in GraphPad Prism 9 and are described in figure legends where applicable. When 2 groups were compared, a two-tailed Student’s t-test was performed. When more than 2 groups were compared, an ordinary one-way ANOVA with Dunnett’s multiple comparisons test was performed. Asterisks denote *p*-value range, where *, *p* < 0.05; **, *p* < 0.01; ***, *p* < 0.001; and ****, *p* < 0.0001.

## ACKNOWLEDGMENTS

The authors would like to thank Destiny Simpson and William Rodgers for helpful discussions and manuscript editing, Franklin Hays for advice with molecular dynamics simulations, and Hunter Porter on assistance with NGS data analysis.

