## Supplementary Information for "High Throughput Characterization of V(D)J Recombination Signal Sequences Redefines the Consensus Sequence"

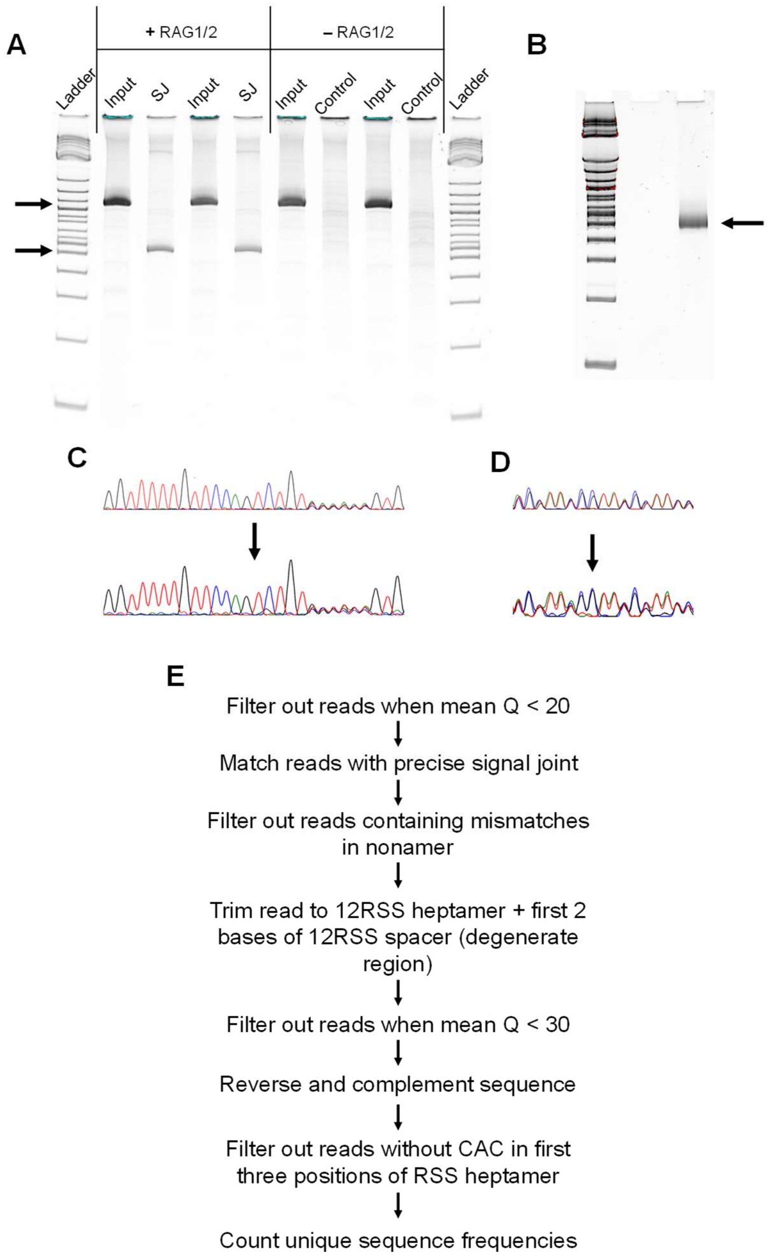


**Fig. S1.** SARP-seq control experiments **(A)** Selective PCR amplification of consensus RSS in pSARP-12R4-9 parent construct (pMX-INV) with and without RAG1/2 co-expression. Arrows indicate expected molecular weights for the input PCR product (higher arrow), which amplifies regardless of V(D)J recombination, and the signal joint PCR product which amplifies V(D)J recombination signal joints. **(B)** SARP-seq output library subjected to NGS. **(C)** Sanger sequencing electropherogram of partially degenerate 12-RSS that was subjected to the SARP-seq protocol *without* RAG1/2 expression **(D)** Sanger sequencing electropherogram of partially degenerate DNA sequence incorporated upstream of pSARP-12R4-9 and subjected to the SARP-seq protocol without RAG1/2 expression. The added degenerate DNA was used to further test whether the SARP-seq protocol skews base degeneracy. **(E)** Flow chart describing SARP-seq NGS data analysis pipeline.

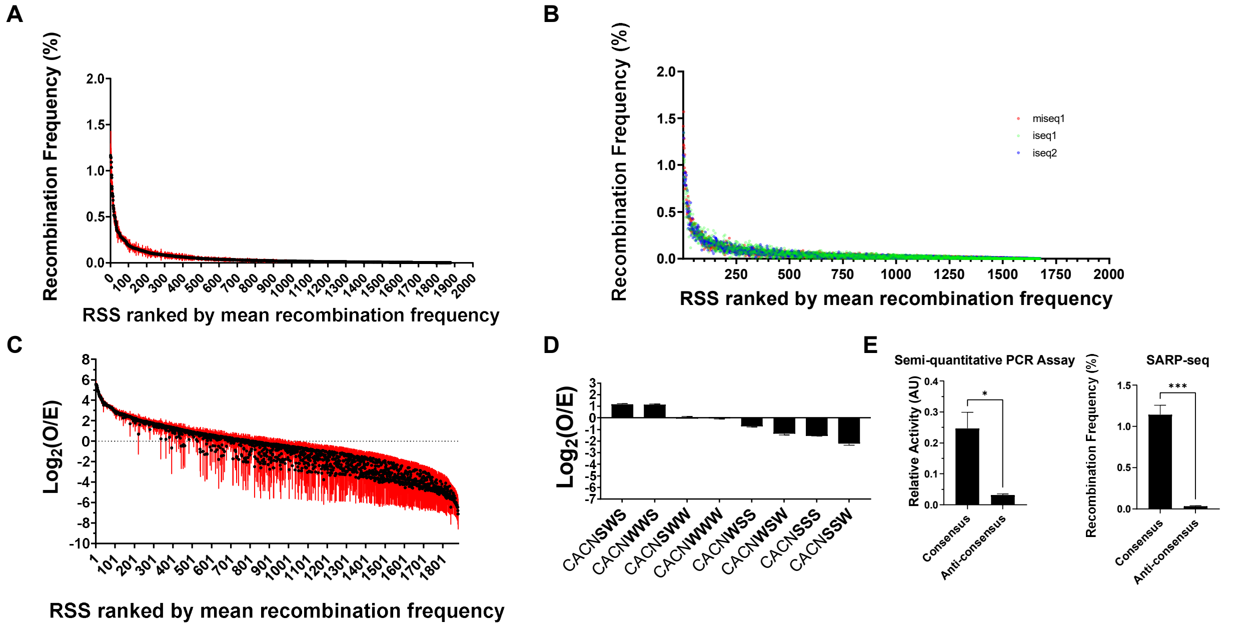


**Fig. S2.** RSS recombination frequency analysis **(A)** RSS recombination frequencies plotted with SEM of n = 3 replicates. **(B)** RSS recombination frequencies with minimum Q30 filtering plotted with SEM of n = 3 replicates. **(C)** RSS recombination frequencies expressed as $\log_2$(O/E) where *O* is the observed recombination frequency of an RSS and *E* is the expected recombination frequency of the same RSS if VDJ recombination was random and non-specific. **(D)** Recombination frequencies for respective RSS motifs where W = A/T and S = C/G. **(E)** Semiquantitative PCR assay (left) and SARP-seq assay (right) comparing consensus RSS (CACAGTGAT) recombination to anti-consensus RSS recombination (CACGTACAT). Bar chart depicts mean and SEM for n = 3 replicates. Consensus and anti-consensus recombination was compared with a two-tailed Student's t-test (*, $p < 0.05$; ***, $p < 0.001$)

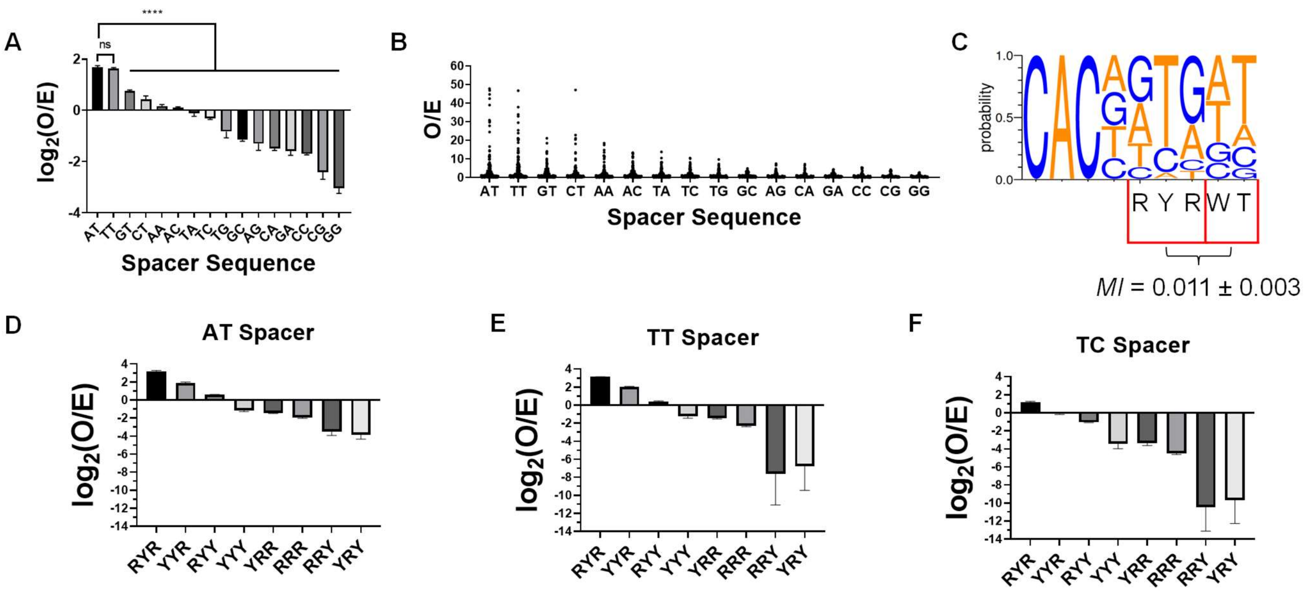


**Fig. S3.** Recombination frequencies for various RSS spacer sequences. **(A)** Mean recombination frequency for each RSS spacer sequence with SD for n = 3 replicates. *p*-values were calculated with an ordinary one-way ANOVA with Dunnett's multiple comparisons test (**** denotes $p < 0.0001$, ns denotes $p > 0.05$). **(B)** Dot plots depicting mean recombination frequencies (n = 3) of each RSS heptamer sequence with corresponding spacer sequence on x-axis. **(C)** Mutual information ± SD shared between R/Y motifs for heptamer positions H5-H7 and all RSS spacer sequences (n = 3). Information content is expressed in nats and calculated using the equation $MI = H_h + H_s - H_{hs}$ where $MI$ is the mutual information shared between the spacer and heptamer R/Y motifs, $H$ is the entropy of RSS heptamer R/Y motifs ($H_h$), spacer sequence motifs ($H_s$), or joint heptamer-spacer sequence motifs ($H_{hs}$) calculated using the equation $H = -\sum P ln(P)$ where $P$ is the sequence motif probability (1). **(D-F)** Recombination frequency of each RSS heptamer R/Y motif with a **(D)** AT **(E)** TT or **(F)** TC spacer sequence. Bar charts depict mean and SD for n = 3 replicates.

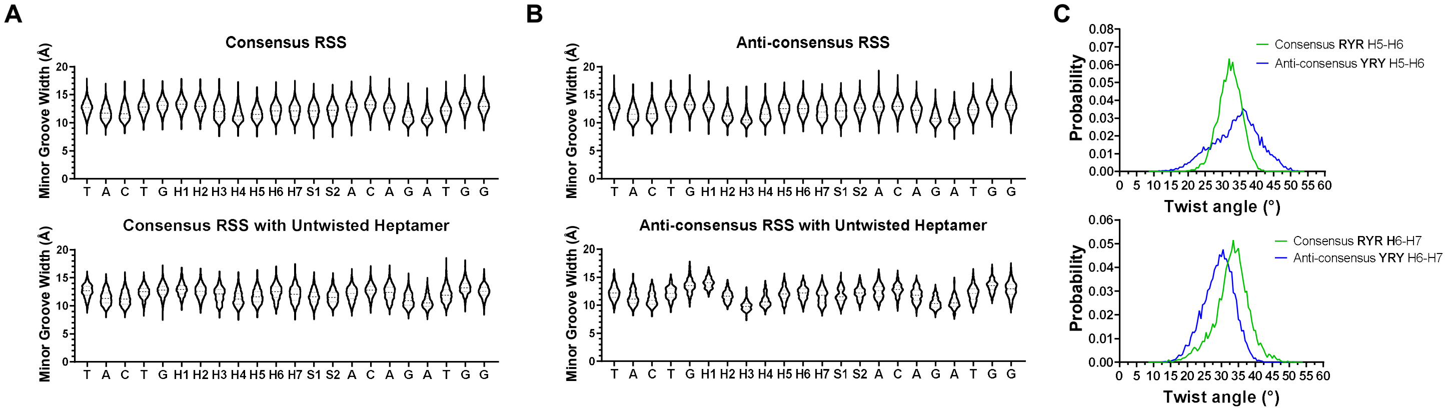


**Fig. S4.** Molecular dynamics simulations of recombination signal sequences. **(A-B)** Top violin plots depict minor groove width measurements for every step of **(A)** consensus and **(B)** anti-consensus simulations. Bottom violin plots depict minor groove width measurements when base-pair step H1-H2 was untwisted (twist angle < 17°). Minor groove widths were measured as inter-phosphate distances using X3DNA "analyze." **(C)** Twist angle probability distributions for base-pair steps H5-H6 (top) and H6-H7 (bottom).

**Table S1.** SARP-seq primer list

| | |
|---|---|
| 12RSS4-9 | GTCTACGAACGAATTCCTACTGCACNNNNNNACAGACTGGAACAAAAACCAGATCGG AAGAGCGTCGTGTAGGGAAAGAGTGTAGATCTCGGTGGTCGCCGTATCATT |
| Duplexing primer | CAGTTCAGGTACGCGTAATGATACGGCGACCACCGAGATCT |
| Nest FWD | TCAAAGTAGACGGCATCGCAG |
| Nest RVS | TCGTCCTTGAAGAAGATGC |
| Input RVS | GAATGCTCGTCAAGAAGACAG |
| P7 primer | AATGATACGGCGACCACCGAGATCT |
| iSeq 1 P5 primer | CAAGCAGAAGACGGCATACGAGATTGGACGTAGCCTTCGGGCATGG |
| miSeq P5 primer 1 | GTGACTGGAGTTCAGACGTGTGCTCTTCCGATCTTGGACGTAGCCTTCGGGCATGG |
| miSeq P5 primer 2 | CAAGCAGAAGACGGCATACGAGATTAAGGCGAGTGACTGGAGTTCAGACGTGTGCTC |

**Movie S1 (separate file).** Movie showing RAG1 bound to minor groove of RSS heptamer. Movie was made using public data from a previous study (PDBID 6OEM) (2). White labels indicate heptamer positions H4 and H7. The RSS heptamer is colored red. RAG1 is represented as ribbon model with stick model of sidechains contacting the RSS minor groove.

**Dataset S1 (separate file).** SARP-seq read counts ranked by normalized mean recombination frequency. Read count data was used to generate all SARP-seq figures for this manuscript.

**Dataset S2 (separate file).** SARP-seq data compared to publicly available END-seq data (3) for *Tcra* $J_{\alpha}$-genes with 25 or more END-seq counts and an RSS heptamer sequence beginning with CAC. To account for position-specific effects on *Tcra* gene recombination, END-seq reads for each $J_{\alpha}$-gene were quantified relative to reads corresponding to 2 adjacent $J_{\alpha}$-genes. For example, $J_{\alpha}$-50 normalized count value was calculated from *O* observed reads divided by *E* which is the mean read count for $J_{\alpha}$-53, $J_{\alpha}$-52, $J_{\alpha}$-50, $J_{\alpha}$-49, and $J_{\alpha}$-48.
